# Azole resistance associated regulatory motifs within the promoter of *cyp51A* in *Aspergillus fumigatus*

**DOI:** 10.1101/2022.02.07.479494

**Authors:** Alexander Kühbacher, Mandy Peiffer, Peter Hortschansky, Petra Merschak, Michael J Bromley, Hubertus Haas, Axel A Brakhage, Fabio Gsaller

## Abstract

*Aspergillus fumigatus* is one of the deadliest fungal species causing hundreds of thousands of deaths each year. As azoles provide the preferred first-line option for treatment of Aspergillosis, the increase in rates of resistance and the poor therapeutic outcomes for those infected with a resistant isolate constitutes a serious global health threat. Azole resistance is frequently associated with specific tandem repeat duplications of a promoter element upstream of *cyp51A*, the gene which encodes the target for this drug class in *A. fumigatus*. This promoter element is recognized by the activating transcription factors SrbA and AtrR. This region also provides a docking platform for the CCAAT-binding-complex (CBC) and HapX that cooperate in the regulation of genes involved in iron-consuming pathways including *cyp51A*. Here, we studied the regulatory contribution of SrbA, AtrR, CBC and HapX binding sites on *cyp51A* expression and azole resistance during different iron availability employing promoter mutational analysis and protein/DNA interaction analysis. This strategy revealed iron status-dependent and -independent roles of these regulatory elements. We show that promoter occupation by both AtrR and SrbA is required for iron-independent steady-state transcriptional activation of *cyp51A* and its induction during short-term iron exposure relies on HapX binding. We further uncover the HapX binding site as repressor element the disruption of which elevates *cyp51A* expression and azole resistance regardless of iron availability.

## Importance

First-line treatment of aspergillosis typically involves the use of azole antifungals. Worryingly, their future clinical use is challenged by an alarming increase in resistance. Therapeutic outcomes for these patients are poor due to delays in switching to alternative treatments and reduced efficacy of salvage therapeutics. Our lack of understanding of the molecular mechanisms that underpin resistance hampers our ability to develop novel therapeutic interventions. In this work, we dissect the regulatory motifs associated with azole resistance in the promoter of the gene that encodes the azole drug target Cyp51A. These motifs included binding platforms for SrbA and AtrR as well as the CCAAT-binding-complex and HapX. Employing mutational analyses, we uncovered crucial *cyp51A* activating and repressing functions of their binding sites. Remarkably, disrupting binding of the iron regulator HapX increased *cyp51A* expression and azole resistance in an iron-independent manner.

## Observation

The human mold pathogen *Aspergillus fumigatus* is the most common cause of invasive aspergillosis. Azole antifungals constitute the major drug class employed for first-line treatment of this life-threatening disease (1). Over the past years a worrying increase in resistance to this drug class has been reported and the *Centers for Disease Control and Prevention* in the United States have recently added azole resistant *A. fumigatus* on their watch list for antibiotic resistance threats (2).

Azoles target the ergosterol biosynthesis enzyme Cyp51, which catalyzes 14α-demethylation of eburicol in *A. fumigatus* (3). Cyp51 inhibition decreases ergosterol production and causes eburicol accumulation, which in turn leads to increased production of toxic C14-methylated sterols (4). Although

*A. fumigatus* expresses two Cyp51 isoenzymes (Cyp51A and Cyp51B), azole resistance mechanisms are rarely linked to mutations of *cyp51B* (5, 6). Two predominant mechanisms of resistance, designated TR34/L98H and TR46/Y121F/T298A, both harbor a tandem repeat (TR) within the promoter of *cyp51A* (*Pcyp51A*) coupled with mutations in the coding sequence leading to amino acid alterations in the enzyme (7, 8). *In vitro* studies using recombinant strains that were transformed with *cyp51A* mutant alleles demonstrated that the TRs elevate *cyp51A* expression, which results in increased azole resistance (9, 10). To date several TR variants have been discovered in azole resistant clinical isolates including the aforementioned TR34 and TR46 as well as TR53 and TR120 (7, 8, 11, 12). It is important to note that the 34mer (wt34) duplicated in TR34 is also present in all the other TRs, suggesting this DNA region as key element driving azole resistance (13, 14). Previous work suggested that wt34, displayed in Fig. 1A, contains binding sites for the *cyp51A* activating factors SrbA and AtrR (13, 15, 16). Overlapping the AtrR binding site, wt34 harbors an imperfect CCAAT box (5’-CGAAT-3’) recognized by the CCAAT-binding-complex (CBC) (13, 17). In close proximity thereof, a HapX binding motif is located (13). Depending on the cellular iron status, the CBC and HapX cooperatively (CBC:HapX) activate or repress the expression of genes involved in iron-dependent pathways such as *cyp51A* (18-20). In addition to *cyp51A*, AtrR and SrbA as well as the CBC:HapX were shown to directly regulate the expression of numerous other target genes including those coding for metabolic enzymes as well as transcriptional regulators (13, 15, 16, 21-23). Hence, it is possible that changes in individual gene expression, resulting from inactivation of these regulators, occur indirect. In this study, we generated mutants harboring disrupted binding motifs of each SrbA, AtrR, the CBC and HapX to study their direct contribution to *cyp51A* regulation and azole resistance. Due to the involvement of these regulators in the control of iron metabolism (15, 21, 23, 24), we monitored the effects of disrupted binding sites on *cyp51A* expression during different iron abundance.

**Fig. 1.**
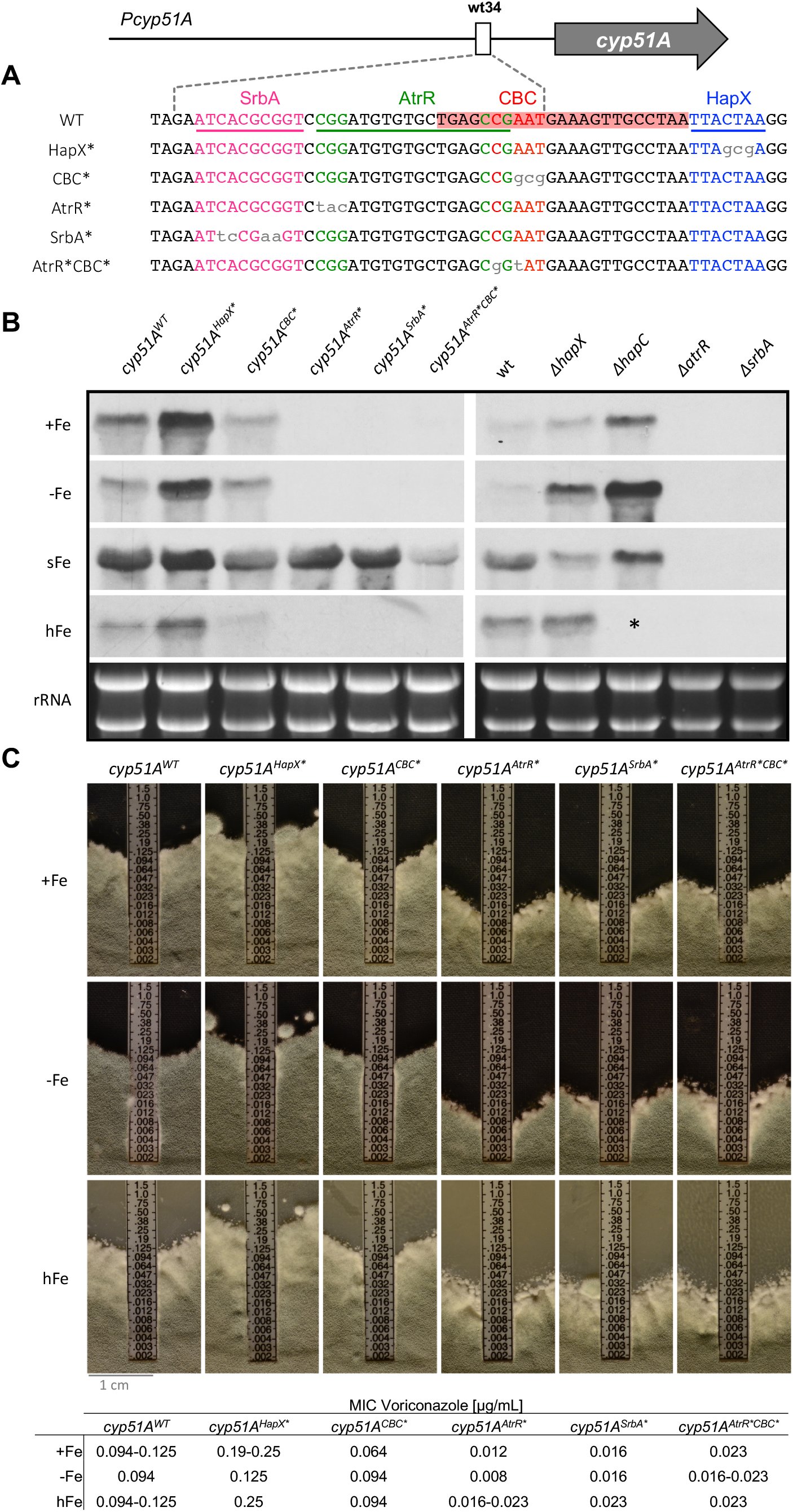
*cyp51A* expression analysis and voriconazole susceptibility testing of *Pcyp51A* mutants. (A) To disrupt interaction of AtrR, SrbA, the CBC and HapX at the 34mer, specific base-pair changes (grey, small letters) were introduced into their binding motifs. Binding sites of SrbA, AtrR and HapX are underlined, the DNA area covered by the CBC is highlighted in red. (B) *cyp51A* transcript levels were determined in mutants containing *cyp51A* expression constructs under control of differently mutated *Pcyp51A* as well as the corresponding transcription factor loss of function mutants. (C) Voriconazole susceptibility of *Pcyp51A* mutants was analyzed using E-test. *Δ*hapC* was not able to grow in the presence of hFe.

Prior to *in vivo* expression analyses, the impact of the introduced mutations on the binding affinity of these transcription factors was analyzed *in vitro* using surface plasmon resonance (SPR) protein-DNA interaction analysis. The binding affinity of SrbA and AtrR was severely affected by the introduced mutations in their specific predicted binding sites (Fig. 2; AtrR: F&H, panel 5; SrbA: I, panel 6) but largely unaffected by mutations in the adjacently located putative binding motifs of the other transcription factors (Fig. 2; panel 5 for AtrR, panel 6 for SrbA). This is particularly important to discriminate binding of the CBC and AtrR as their binding motifs overlap (Fig. 1A). Comparing DNA binding affinities of the CBC (Fig. 2A/panel 1), with that of premixed CBC and HapX (Fig. 2A/panel 4) and HapX to preformed CBC-DNA (Fig. 2A/panel 2 or 3) indicated that HapX improves binding of the CBC to DNA. This was even true for its interaction at the mutated CBC binding motif, which was not recognized by the CBC alone (Fig. 2C-D/panel 1 and 4). The observed assistance of HapX for CBC-DNA binding might be due to the fact that the CBC recognition motif is an imperfect CCAAT box with only moderate CBC affinity (21). Mutation of the predicted HapX binding site impaired binding of HapX to preformed CBC-DNA complexes (Fig. 2B/panel 2 and 3) as well as that of premixed CBC and HapX (Fig. 2B/panel 4), which is consistent with the idea that HapX assists CBC-DNA interaction at wt34.

**Fig. 2.**
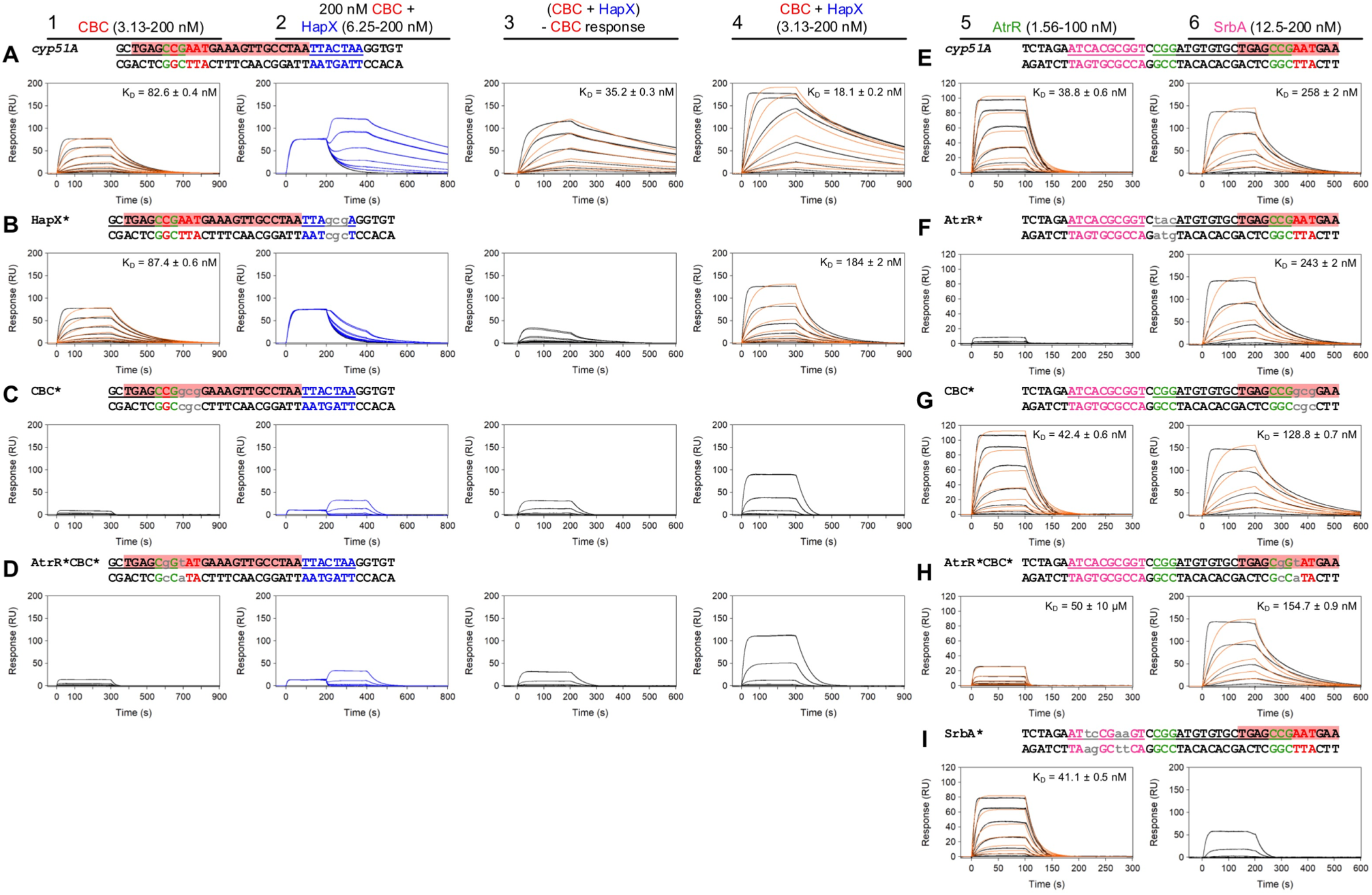
SPR-based binding analysis of SrbA, AtrR, CBC and HapX to their consensus binding motifs at the 34mer. Sensorgrams for binding of CBC to DNA (1), HapX to preformed CBC-DNA complexes (2) premixed CBC and HapX to DNA (4), AtrR to DNA (5) and SrbA to DNA (6). Sensorgrams in panel 3 show the association/dissociation responses of HapX on preformed CBC-DNA. Hereby, CBC response was subtracted (co-injection of buffer) from HapX co-injection responses. Interaction with non-mutated (A and E) as well as mutated binding sites (B-D and F-I) was monitored. Binding responses of the indicated SrbA, AtrR, CBC or HapX concentrations injected in duplicate (black lines) are overlaid with the best fit derived from a 1:1 interaction model, including a mass transport term (red lines). Binding responses of CBC-DNA-HapX ternary complex formation (panel 2, blue lines) were obtained by concentration-dependent co-injection of HapX on preformed binary CBC-DNA complexes after 200 s within the steady-state phase. Dissociation constants (K_D_) of the complexes are displayed inside the graphs.

For *in vivo* expression analyses, *cyp51A* gene cassettes comprising non-mutated and mutated *Pcyp51A* variants (Fig. 1A), analogous to those assessed by SPR (Fig. 2), were integrated at its native locus (Fig. S1). Differential expression resulting from disrupted binding sites was determined by comparing *cyp51A* transcript levels of *Pcyp51A* mutants (*cyp51A*^*AtrR\**^, *cyp51A*^*SrbA\**^, *cyp51A*^*CBC\**^, *cyp51A*^*HapX\**^ and *cyp51A*^*AtrR*CBC\**^) to those of the strain carrying the non-mutated promoter (*cyp51A*^*WT*^). To identify potential iron status-specific effects, *cyp51A* expression was monitored during iron starvation (-Fe, 0 mM FeSO_4_), sufficiency (+Fe, 0.03 mM FeSO_4_), excess (hFe, 5 mM FeSO_4_) and during short-term iron adaptation by supplementing -Fe cultures for 30 min with iron (sFe). Furthermore, the regulation pattern of *Pcyp51A* mutants was compared to those of the respective transcription factor loss of function mutants.

HapX, together with the CBC, was previously shown to mediate repression of *cyp51A* during iron starvation and upregulation during short-term iron exposure (19). This is in accordance with the *cyp51A* transcript levels found in wt and mutant strains lacking HapX (Δ*hapX*) or CBC (Δ*hapC*) as well as in the strains carrying the unmutated (*cyp51A*^*WT*^) and mutated (*cyp51A*^*HapX\**^) HapX binding motif, respectively (Fig. 1B; importantly, sFe mediated induction is characterized by increased transcript levels in sFe compared to -Fe). Interestingly, in contrast to HapX inactivation (Δ*hapX*), mutation of its binding site (*cyp51A*^*HapX\**^) caused increased *cyp51A* expression also during iron sufficiency and iron excess. Remarkably, mutation of the CBC binding site (*cyp51A*^*CBC\**^) resulted in lower *cyp51A* transcript levels when compared to mutation of the HapX binding site (*cyp51A*^*HapX\**^) and, moreover, did not abolish *cyp51A* induction during sFe (Fig. 1B). Therefore, we hypothesize that HapX dominates the sFe induction of *cyp51A* and DNA-binding of the CBC plays a minor role for iron regulation in this case, which could be related to the potential stabilizing properties of HapX found for *in vitro* binding of the CBC to DNA (Fig. 2A,C,D/panels 1, 4). The different expression pattern observed in *cyp51A*^*HapX\**^ and *cyp51A*^*CBC\**^ compared to their corresponding loss of function mutants could be a result of indirect regulatory defects or, the potential stabilization of the CBC or HapX by other proteins at the 34mer.

Mutation of either the AtrR or the SrbA binding motif (*cyp51A*^*AtrR\**^ and *cyp51A*^*SrbA\**^) as well as loss of the corresponding transcription factors (Δ*atrR* and Δ*srbA*) led to severely decreased *cyp51A* expression during -Fe, +Fe as well as hFe (Fig. 2B), which agrees with the *cyp51A* activating role of AtrR and SrbA. These findings suggest that the AtrR and SrbA motifs analyzed here are the physiologically most relevant *in vivo* binding sites despite the presence of further predicted SrbA binding motifs within *Pcyp51A* (13, 24, 25). The results further emphasize the interdependency of these two transcription factors as previously shown for a subset of common target genes (15, 26), and demonstrate for first time that both SrbA and AtrR have to bind to wt34 in *Pcyp51A* for steady-state transcriptional activation. Surprisingly, in contrast to Δ*srbA* and Δ*atrR, cyp51A*^*SrbA\**^ and *cyp51A*^*AtrR\**^ were still able to fully induce *cyp51A* expression during sFe. These results demonstrate that AtrR and SrbA are both required for sFe induction but not their binding to wt34. SrbA has previously been shown to be required for transcriptional activation of HapX (24). Consequently, SrbA and AtrR might be indirectly required for sFe induction, which is presumably exclusively mediated by the CBC-HapX complex.

The mutation impairing binding of both AtrR and CBC by affecting the overlapping consensus motifs (*cyp51A*^*AtrR*CBC\**^, Fig. 1A) displayed a combination of the defects caused by the individual disruption of AtrR (*cyp51A*^*AtrR\**^) and CBC (*cyp51A*^*CBC\**^) binding, i.e. *cyp51A* transcript levels were severely reduced during -Fe, +Fe and hFe and sFe induction was significantly diminished (Fig. 1B).

In the next step, we monitored the impact of disrupted binding motifs on iron-dependent voriconazole susceptibility using E-test (Biomerieux®). Mutants with defective steady-state *cyp51A* activation (*cyp51A*^*AtrR\**^, *cyp51A*^*SrbA\**^ and *cyp51A*^*AtrR*CBC\**^) showed increased susceptibility during -Fe, +Fe and hFe (Fig. 1C). These data clearly demonstrate that defective binding of either AtrR or SrbA to *Pcyp51A* at wt34 causes dramatically increased azole susceptibility. The resistance pattern of *cyp51A*^*CBC\**^ was similar to that of *cyp51A*^*WT*^ during -Fe and hFe. In agreement with reduced *cyp51A* expression in *cyp51A*^*CBC\**^, this strain was slightly more susceptible to voriconazole at +Fe. *cyp51A*^*HapX\**^, showing increased steady-state *cyp51A* transcript levels during different iron abundance, displayed elevated resistance independent of the iron status.

## Conclusions

TR-based *cyp51A* overexpression represents a major cause driving azole resistance in *A. fumigatus* (7, 8, 11, 12). Hence, a profound understanding of regulatory mechanisms associated with TRs is crucial to elucidate therapeutic strategies that counteract resistance. In this work we explored iron status-dependent, regulatory functions of AtrR, SrbA, CBC and HapX binding motifs located at the azole resistance associated 34mer, the key enhancer element involved in TR-based *cyp51A* upregulation (13). We identified iron-independent, *cyp51A*-activating functions of AtrR and SrbA binding sites during steady-state cultivation and found the HapX binding motif to be crucial for the induction of the gene during short-term iron exposure. Moreover, despite the known -Fe specific repressing role of HapX on iron consuming genes such as *cyp51A* (18, 19, 23), our results suggest the requirement of its functional binding motif for the repression of *cyp51A* during steady-state growth, regardless of the iron availability. Hampering this repression by disrupting the HapX binding site elevates resistance to voriconazole and, thus, unveils its binding motif as potential azole resistance hotspot.

## Materials and Methods

For details see Supplemental Materials and Methods.

